# The Interaction of Viperin with Hepatitis C Virus Non-Structural Protein 5A Inhibits the Catalytic Activity of Viperin

**DOI:** 10.1101/824458

**Authors:** Soumi Ghosh, Ayesha M. Patel, Timothy J. Grunkemeyer, Arti B. Dumbrepatil, Kelcie Zegalia, Robert T. Kennedy, E. Neil G. Marsh

## Abstract

The radical SAM enzyme viperin exerts a wide range of antiviral effects through both the synthesis of the antiviral nucleotide 3’-deoxy-3’, 4’-didehydro-CTP (ddhCTP) and through its interactions with various cellular and viral proteins. Here we investigate the interaction of viperin with hepatitis C virus non-structural protein 5A (NS5A) and the host sterol regulatory protein, vesicle-associated membrane protein A (VAP-33). NS5A and VAP-33 form part of the viral replication complex that is essential for copying the RNA genome of the virus. Using transfected enzymes in HEK293T cells, we show that viperin binds to both NS5A and VAP-33 independently and that this interaction is dependent on all three proteins being localized to the ER membrane. Co-expression of viperin with VAP-33 and NS5A was found to reduce NS5A levels, most likely by increasing the rate of proteasomal degradation. However, co-expression of viperin with VAP-33 and NS5A also reduces the specific activity of viperin by ~ 3-fold. This observation suggests that NS5A may have evolved to bind viperin as a strategy to reduce ddhCTP synthesis, thereby reducing possibility of the replication complex introducing this chain-terminating nucleotide during genome synthesis.

## Introduction

Viral infection of eukaryotic cells initiates a number of viral recognition strategies aimed at preventing the spread of infection. Induction of antiviral cytokine interferons (IFN) is one of the primary innate immune defences against viruses. Type I IFN induces the expression of numerous interferon-stimulated genes (ISGs) that limit viral replication. Viperin (Virus inhibitory protein, Endoplasmic Reticulum-associated, Interferon-inducible) is one of the ISGs that have been implicated directly as having antiviral activity. Viperin appears to exert antiviral activity against a wide range of viruses by and has been show to interact with various cellular and viral proteins. Especially, it has been shown to inhibit several flaviviridae family viruses; including Hepatitis C Virus (HCV) (1–3), dengue virus (DENV)(4,5), tick-borne encephalitis virus (TBEV)(6–8), West Nile virus (WNV)(4), and Zika virus (ZIKV)(8–11).

The molecular basis of viperin’s inhibitory effects seems to be virus-specific. For HCV and DENV, it was shown to inhibit viral replication by interacting with the non-structural proteins(1,5,11,12), while for TBEV, it impedes the synthesis of positive-strand viral RNA(7). A recent study revealed that viperin can also inhibit ZIKV and TBEV replication by interacting with both structural and non-structural viral proteins(8). The regulation of viral proteins by viperin unveils the mechanistic insights of its antiviral activity. However, the underlying unifying mechanism by which viperin targets a broad range of viruses is still under investigation.

Viperin is a radical S-adenosyl-L-methionine (SAM)-dependent enzyme (13–15), a superfamily of enzymes that catalyses a wide range of reactions involving free radical intermediates. Of the more than 110,000 radical SAM enzymes, viperin is one of only eight found in humans (16). The superfamily is characterized structurally by a partial (βα)_6_ barrel fold with a CxxxCxxC motif [*Figure 1(a)*]; the cysteinyl residues of which chelate the iron-sulfur [Fe_4_S_4_] cluster that is essential for radical generation. A 5’-deoxyadenosyl radical (5’-dAdo•) is generated from SAM by 1-electron reduction mediated by the iron-sulfur cluster. The adenosyl radical then abstracts a hydrogen atom from the substrate to form 5’-deoxyadenosine (5’-dAdo) as a by-product [*Figure 1(c)*]. In subsequent steps, the substrate-radical intermediate can undergo a diverse array of chemical transformations, depending upon the enzyme (17–19).

**Figure 1:**
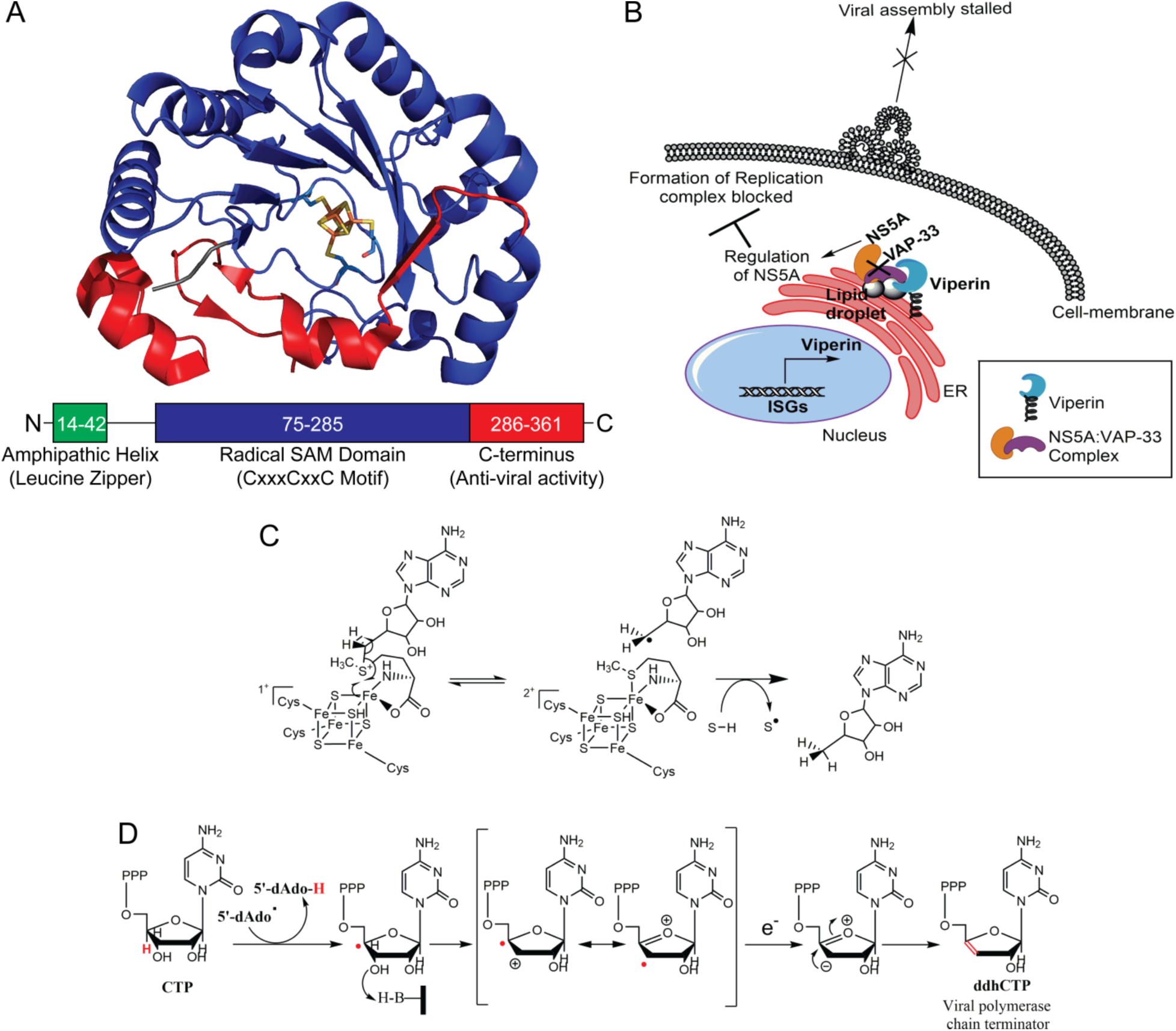
Overview of viperin’s function in synthesizing the antiviral nucleotide ddhCTP and inhibiting Hepatitis C virus replication. (a) Crystal structure of mouse-viperin (PDB ID: 5VSL), showing the three major domains, including the [Fe_4_S_4_] cluster-chelating CxxxCxxC motif. (b) Role of viperin in restricting HCV replication complex formation by interacting with the HCV protein Non-structural 5A (NS5A) through vesicle-trafficking host protein VAP-33. (c) Mechanism of one-electron reductive cleavage of SAM by the [Fe_4_S_4_] cluster in radical SAM enzymes to generate 5’-deoxyadenosyl radicals. (d) Proposed mechanism for the radical-mediated dehydration of CTP to form the antiviral nucleotide 3’-deoxy-3’, 4’-didehydro-CTP (ddhCTP).

The enzymatic activity of viperin was only recently elucidated; it was shown to catalyse the formation of a novel antiviral ribonucleotide by dehydration of CTP to 3’-deoxy-3’, 4’-didehydro-CTP (ddhCTP) [*Figure 1(d)*]. ddhCTP acts as a chain terminator for the RNA-dependent RNA polymerases (RdRp) in flaviviruses, thereby explaining one facet of viperin’s antiviral activity (20). However, ddhCTP did not seem to be effective in restricting RdRps from other human viral families. Moreover, viperin’s catalytic activity does not explain viperin’s ability to regulate different metabolic pathways through its interactions with different cellular and viral proteins. For example, viperin was recently shown to modulate immune signalling through the Toll-like receptor-7 and −9 pathways by interactions with interleukin-1 receptor– associated kinase-1 (IRAK1) and E3 ubiquitin ligase TNF receptor–associated factor 6 (TRAF6) (21).

In this study we focus on viperin’s interaction with the Hepatitis C non-structural protein 5A (NS5A), which is an essential component of the viral replication complex (1,3). NS5A localizes at the cytoplasmic face of the endoplasmic reticulum and lipid droplets together with the RdRp, NS5B. Interaction of NS5A with the sterol regulatory host protein, vesicle-associated membrane protein A (VAP-33), is also needed to support HCV replication (12,22). Through FRET analysis (3), confocal microscopy and co-immunoprecipitation studies (1), viperin was shown to interact with NS5A through VAP-33 at lipid droplets. This observation suggests that the interaction between viperin and VAP-33 may interfere with the association of NS5A and VAP-33, thereby perturbing the viral replication complex formation [*Figure 1(b)*]. Mutagenic analysis showed that the C-terminal domain of viperin is required for its interaction with NS5A through VAP-33 in HEK293T cells.

Here we have investigated the interactions between viperin, NS5A and VAP-33 and their effect of virperin’s catalytic activity using proteins transiently expressed in HEK293T cells and with pure proteins *in vitro*. We show that full-length viperin interacts with NS5A in presence of host cell factor VAP-33 and leads to its degradation through the proteasomal degradation pathway. Concomitantly, NS5A and VAP-33 reduce the ddhCTP-forming activity of viperin. The formation of the complex between viperin, NS5A and VAP-33 is dependent on the membrane localization of all three proteins. The present study provides insight into the mechanism of regulation of a viral protein by viperin, coupled with its enzymatic activity.

## Results

### Viperin interacts with NS5A at endoplasmic reticulum

We first examined the interaction of viperin with NS5A and the c-terminal cytoplasmic domain of VAP-33 (hVAPc; human homolog with amino acid residues 156-242), which previous studies (1) had shown is responsible for binding to NS5A. We conducted immunoprecipitation experiments using viperin or viperinΔ3C (viperin variant with iron-sulfur cluster-chelating cysteine residues mutated to alanine) as bait proteins and NS5A and/or hVAPc as prey proteins.

NS5A was found to co-precipitate with viperin *[Figure 2(a)]*, independent of the presence of hVAPc. NS5A was also co-precipitated by the viperinΔ3C mutant *[Figure 2(a)]*, suggesting that the presence of the iron-sulfur cluster is not an essential part of the viperin-NS5A interaction. Both viperin and viperinΔ3C mutant were found to precipitate hVAPc individually. We further confirmed the co-localization of viperin, NS5A and hVAPc by immuno-fluorescence microscopy of fixed HEK293T cells, where NS5A was observed to co-localize with viperin at endoplasmic reticulum in the presence and absence of hVAPc *[Figure SI.1]*

**Figure 2.**
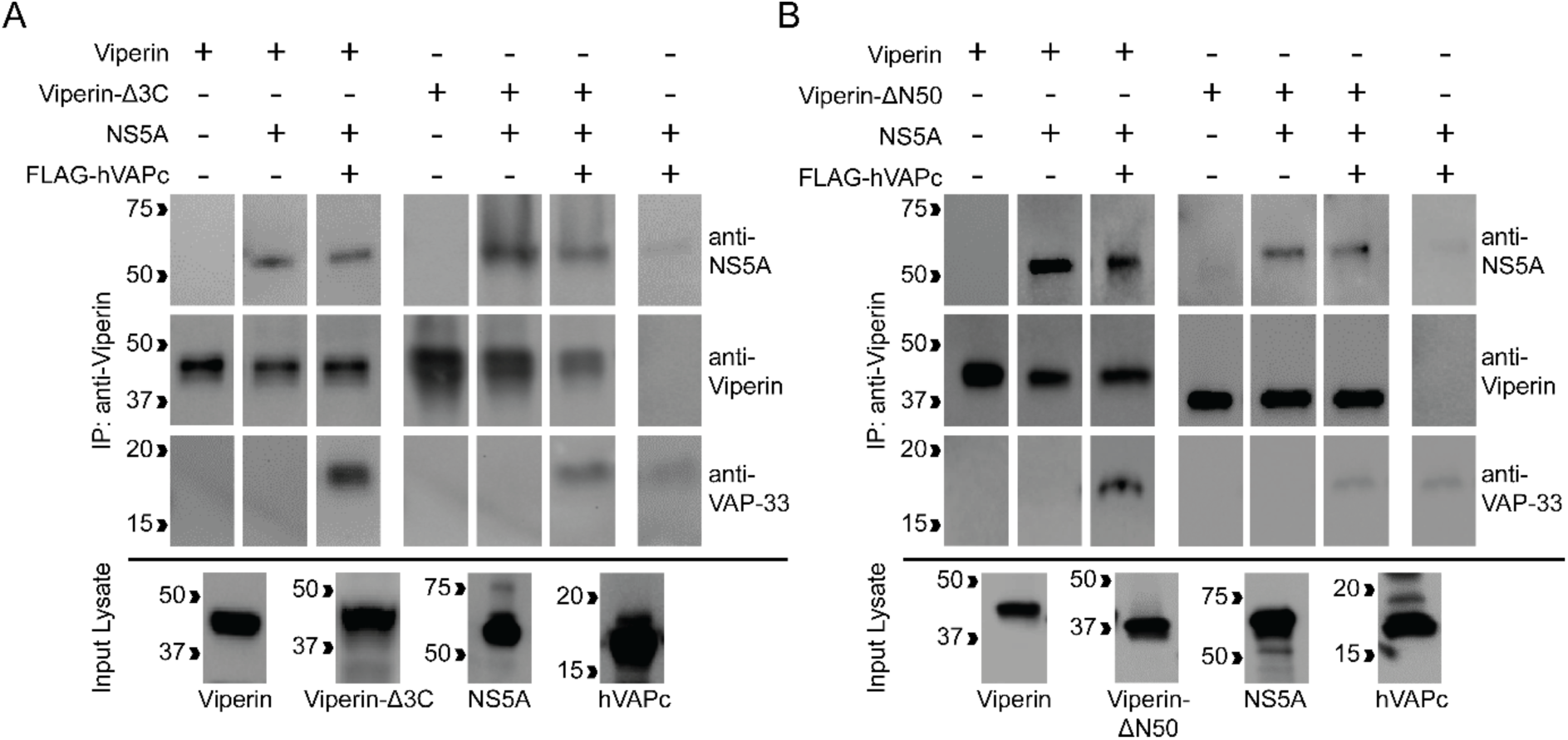
Viperin interacts with NS5A and co-localizes at endoplasmic-reticulum. (a) HEK293T cells transfected viperin or viperin-Δ3C (lacking the iron-sulfur cluster), NS5A and or FLAG-hVAPc (c-terminal of VAP-33) were immunoprecipitated using anti-viperin antibody and blots probed with anti-NS5A monoclonal and anti-FLAG (for hVAPc) antibody. The results demonstrate that viperin and viperin-Δ3C pull down both NS5A and hVAPc and the iron-sulfur cluster is not required for viperin to bind NS5A and hVAPc. (b) Co-immunoprecipitation using viperin or viperin-ΔN50 (lacking the N-terminal amphipathic helix) as bait and NS5A and hVAPc as prey protein. The results demonstrate that the N-terminal amphipathic helix is important for viperin to bind NS5A and hVAPc.

We also assessed the importance of the N-terminal amphipathic helix of viperin for its interaction with NS5A and hVAPc by co-immunoprecipitation using an N-terminal truncated viperin construct (viperin-ΔN50) that lacks the ER localizing amphipathic helix as the bait protein. In this case NS5A and hVAPc showed a weaker interaction with viperin-ΔN50 *[Figure 2(b)]*, indicating that the localization of viperin to the endoplasmic-reticulum is important for its interaction with the target proteins.

### Viperin leads to the degradation of NS5A through proteasomal degradation pathway

Viperin has been found to alter the cellular expression levels of various proteins it interacts with(8,23). Therefore, we next examined how co-expression of viperin altered the cellular levels of NS5A and hVAPc. Proteins were co-transfected into HEK293T cells and after 30 h the cells were harvested and protein expression analysed by immunoblotting. Co-expression with hVAPc had no significant effect on NS5A expression, whereas co-expression with viperin resulted in a small decrease in NS5A levels. Co-expression of both hVAPc and viperin led to the most significant decrease in intra-cellular expression level of NS5A, by ~2-fold, compared to NS5A expressed on its own *[Figure 3(a)]*. No reduction of NS5A levels were observed in control experiments when empty vector (pcDNA3.1) was co-transfected with it. Similar reductions in NS5A levels were observed when NS5A was co-expressed with viperinΔ3C and hVAPc, suggesting the iron-sulfur centre is not required for this effect.

**Figure 3:**
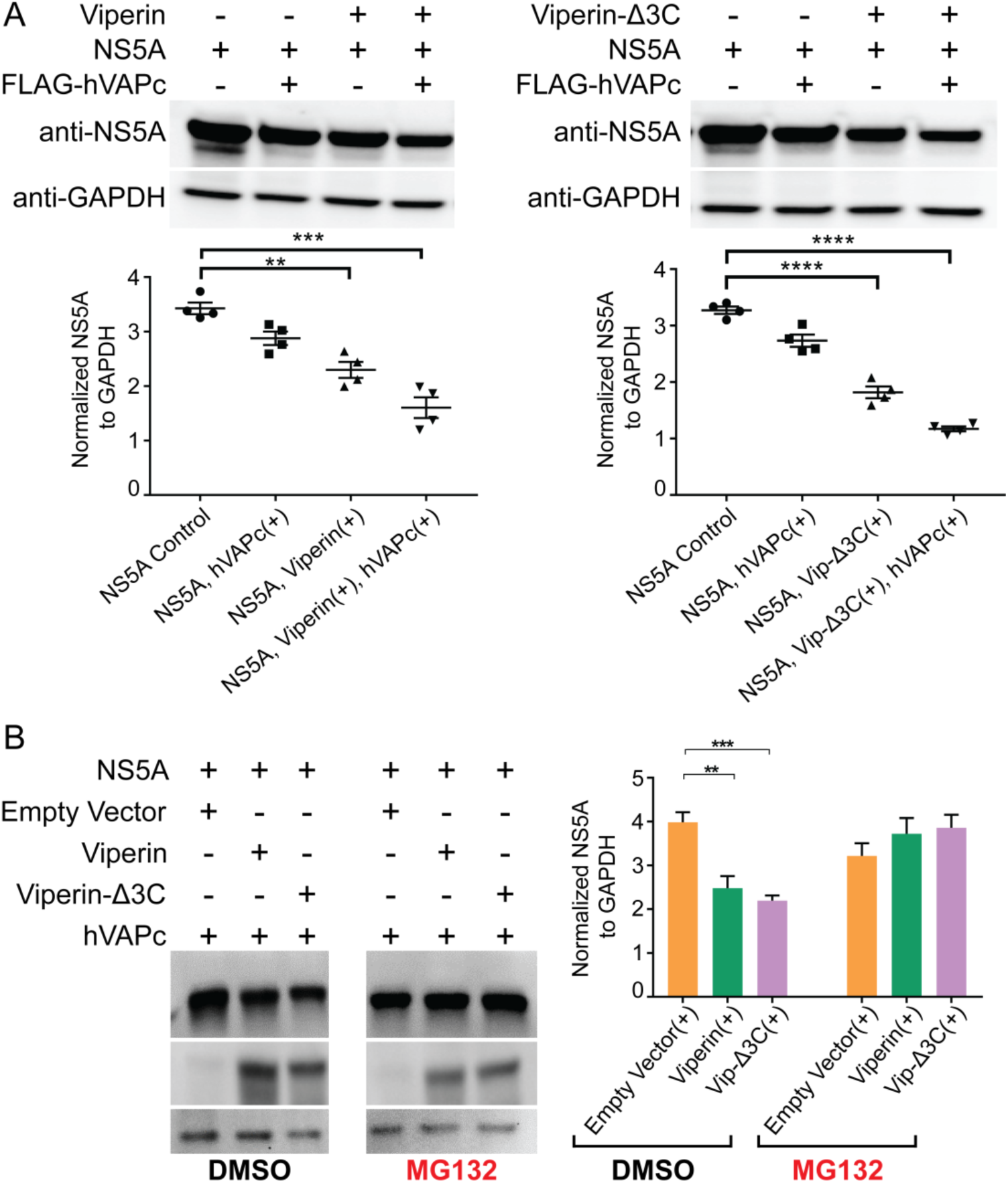
Viperin promotes degradation of NS5A through the proteasomal degradation pathway. (a) HEK293T cells were transfected with NS5A, FLAG-hVAPc and either viperin or viperin-Δ3C. 30 hours post-transfection NS5A levels were visualized by immunoblotting. Co-expression of viperin or viperin-Δ3C and hVAPc significantly decreases NS5A levels (*left panel* *** p = 0.0005, n = 4; *right panel* ****p = 0.0001, n = 4). (b) HEK293T cells were transfected with NS5A, hVAPc and either viperin or viperin-Δ3C. 6 hours post transfection cells were treated with either 50 μM MG-132 (proteasome inhibitor) or DMSO control; 30 hours post-transfection NS5A levels were visualized by immunoblotting. MG-132 reverses the decrease in NS5A levels induced by co-expression of viperin or viperin-Δ3C (** p = 0.0019, n=6; *** p = 0.0002, n=6).

To investigate the reason for the viperin-mediated reduction in NS5A expression, we examined the ability of the proteasomal inhibitor MG-132 to restore NS5A expression levels. The expression levels of NS5A co-expressed with viperin or viperinΔ3C were restored to levels comparable to those of the controls by treatment with MG-132 *[Figure 3(b)]*. These results suggest that viperin promotes the degradation of NS5A through the proteosomal degradation pathway.

### NS5A and hVAPc together repress the reductive SAM cleavage activity of viperin

NS5A is known to antagonize the innate immune system by inactivating various ISGs (24). For example, NS5A was shown to disrupt dimerization and repress the enzymatic activity of Protein kinase R, a critical ISG in cellular antiviral response (25). Therefore, we investigated whether NS5A might alter the catalytic activity of viperin. To examine this possibility, we measured specific activity of viperin in HEK293T cell lysates when co-expressed with NS5A and/or hVAPc. HEK293T cell-lysates were prepared under anaerobic conditions and viperin activity was quantified as described previously (21).

When expressed on its own, viperin showed a specific activity, expressed as a turnover number, k_obs_ = 7.6 ± 0.6 h^−1^. When co-expressed with hVAPc, this activity increased slightly with k_obs_ = 13.1 ± 1.1 h^−1^, whereas co-expression with NS5A did not change the specific activity of viperin, k_obs_ = 8.4 ± 0.7 h^−1^. However, when viperin was co-expressed with both hVAPc and NS5A, the specific activity of viperin was significantly lowered to k_obs_ = 4.0 ±0.4 h^−1^. *[Figure 4, Figure SI.3]*. VAP-33 is a known interaction partner of viperin, therefore, it was not unexpected to see the increase in activity of viperin (about ~1.7-fold relative to single-expressed viperin) in its presence. However, a ~3.3-fold reduction in relative activity of viperin upon addition of NS5A (comparing viperin with hVAPc alone and with both proteins) is quite remarkable; as this deactivation of radical SAM activity of viperin has not been previously observed.

**Figure 4:**
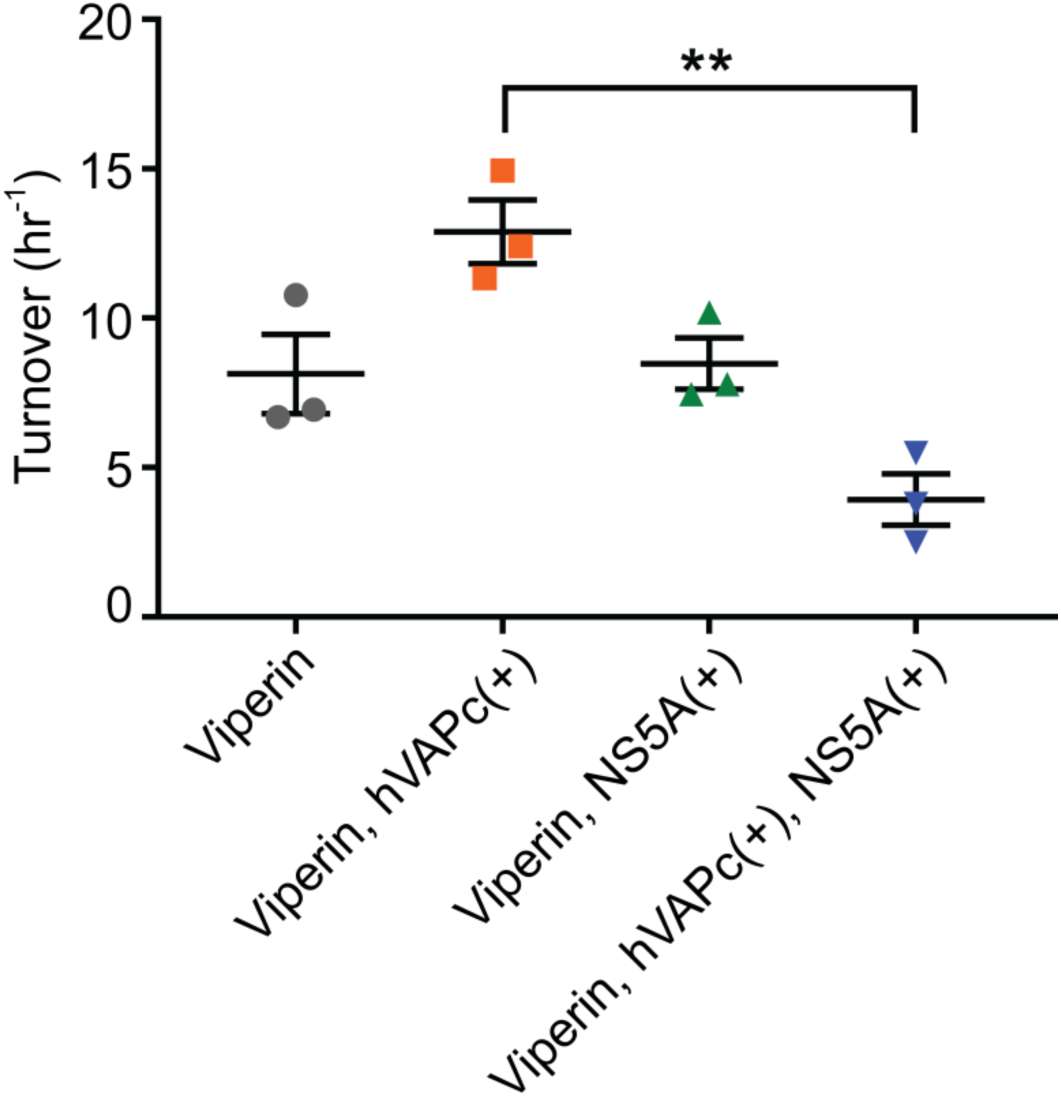
Co-expression of NS5A and hVAPc inhibits reductive SAM cleavage activity of Viperin in HEK293T cell-lysates. (a) Activity of viperin in HEK293T cell-lysates co-expressing either NS5A and/or hVAPc. The amount of 5’-dA produced was determined after 1 hour and normalized to the amount of viperin expressed in the cell extract; data presented as mean ± SEM n = 3. A significant (**p=0.0033) reduction in reductive SAM cleavage activity was observed when viperin was co-expressed with NS5A and hVAPc.

### The membrane-localizing domains of viperin, NS5A and VAP-33 are required for complex formation

To examine the changes in enzymatic activity in more detail we attempted to reconstitute the interactions between viperin, NS5A and hVAPc using purified proteins obtained by over-expression in *E. Coli.* However, this necessitated removing the membrane-binding domains of these proteins so that they could be produced in soluble form. A C-terminal His_6_-tagged viperin lacking the first 50 membrane-associated amino acids (viperin-ΔN50), was purified under anaerobic conditions and the iron-sulfur cluster was reconstituted using procedures as described previously (15). NS5A lacking the first 39 amino acids comprising the membrane-binding amphipathic helix (NS5A-ΔN39), was expressed and purified as a fusion protein with maltose-binding protein (MBP), followed by TEV cleavage and size exclusion chromatography as described in the methods section. The N-terminal His_6_-tagged VAP-33 construct lacking the last 20 amino acids of the trans-membrane domain on the C-terminus (VAP-33-ΔC20), was purchased commercially.

However, when the interactions between these three proteins we examined by immunoprecipitation using an anti-viperin antibody, no protein complexes could be detected *[Figure 5]*. Further experiments designed to detect the formation of protein complexes using the lysine-reactive covalent cross-linking reagent, bis(sulfosuccinimidyl)suberate (BS^3^), similarly failed to detect any inter-protein crosslinking. Consistent with this, the catalytic activity of viperin-ΔN50 was unaffected by the presence of NS5A-Δ39, or VAP-33-ΔC20. Under the conditions of the assay, k_obs_ for viperin-ΔN50 was 4.1 ± 0.3 h^−1^, with the lower activity observed for the truncated construct being consistent with previous observations (21). When mixed with stoichiometric amounts of NS5A-ΔN39 or VAP-33-ΔC20, we measured rates of 2.7 ± 0.2 h^−1^ and 2.8 ± 0.1 h^−1^ respectively. Finally, combining all three proteins yielded a comparable result with k_obs_ = 2.8 ± 0.1 h^−1^ *[Figure 6].* These results suggest that removing the membrane-associated domains from these proteins effectively abolishes the ability of the proteins to form a complex.

**Figure 5:**
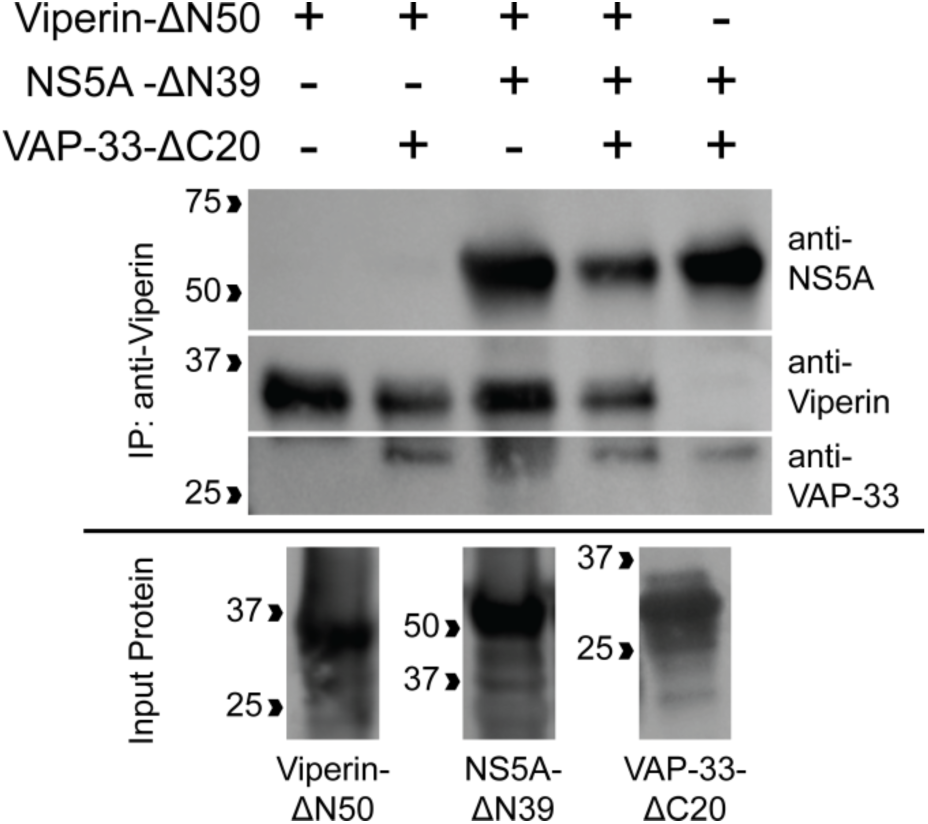
Co-immunoprecipitation of recombinant NS5A-Δ39 and VAP-33-ΔC20 using purified viperin-ΔN50 (lacking the membrane associated sequence) as bait protein *in vitro*: Recombinant viperin was incubated with its possible prey protein recombinant NS5A-ΔN39 and VAP-33-ΔC20, followed by its immunoprecipitation, using anti-viperin antibody. The immunoprecipitated samples were immunoblotted with anti-his antibody to probe for the proteins. No specific interaction was observed between viperin and NS5A in presence and absence of VAP-33, suggesting that the membrane associated sequence of these interacting proteins are important for protein-complex formation.

**Figure 6:**
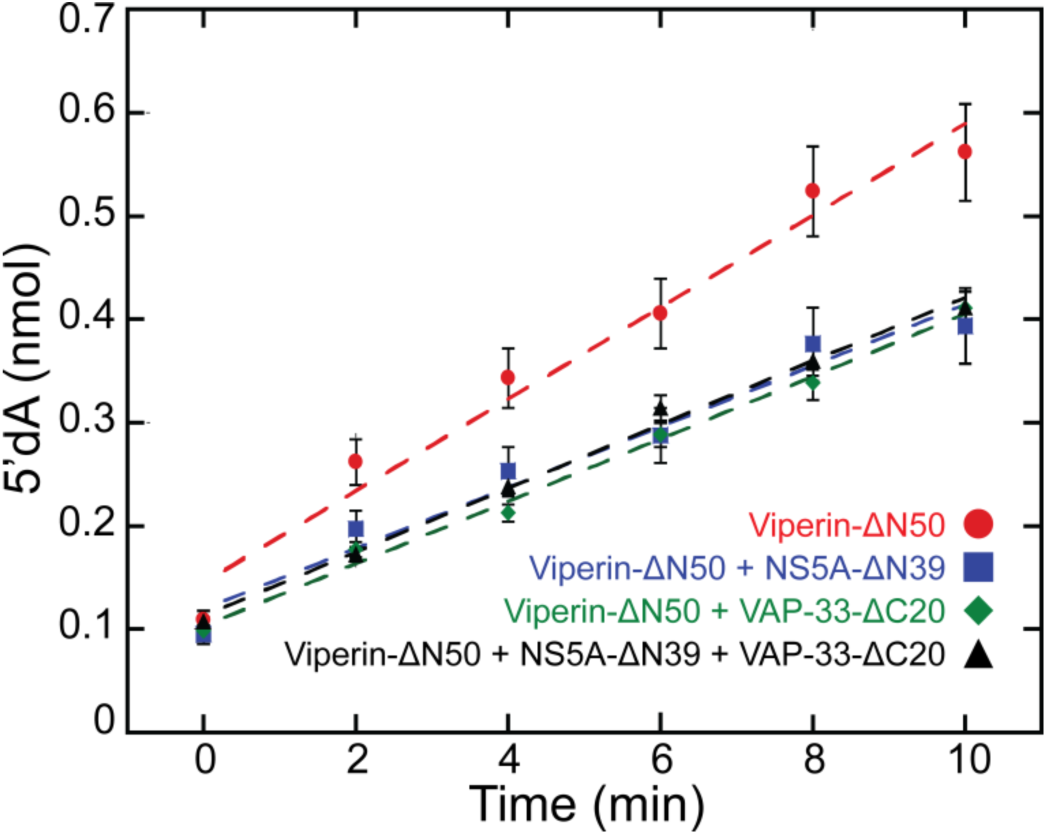
Recombinant NS5A and VAP-33, lacking the membrane associated domain, did not change the reductive SAM cleavage activity of recombinant viperin *in vitro.* Activity of purified viperin (viperin-ΔN50) was assessed by monitoring the rate of 5’-deoxyadenosine production. The turnover of 5’-deoxyadenosine by viperin was calculated from the linear fitting curve. Viperin showed the turnover of 4.1 ± 0.3 h^−1^ alone and 2.7 ± 0.2 h^−1^, 2.8 ± 0.1 h^−1^, 2.8 ± 0.1 h^−1^ when incubated with recombinant NS5A (NS5A-ΔN39), VAP-33 (VAP-33-ΔC20) or both, respectively. Overall, no significant change was observed in the activity of viperin, when combined with purified NS5A and VAP-33.

## Discussion

To date, most of the studies conducted on the antiviral activity of viperin against flaviviridae viruses have been focused on identifying physical interactions between viperin and viral core and non-structural proteins and correlating perturbations of these interactions with changes in viral replication or infectivity. Though the importance of domain-specific interactions between viperin and its target viral proteins has been shown through mutational analysis, no investigation of the effects of these interactions on the recently-revealed catalytic activity of viperin have been undertaken. The discovery that viperin catalyzes the formation of the antiviral nucleotide ddhCTP from CTP raises the question of whether the activity of viperin is modulated by the numerous proteins it has been shown to interact with. Understanding the mechanism by which the synthesis of ddhCTP is viperin regulated by other proteins may open the way to the design of new antiviral therapeutics.

Non-structural protein NS5A from HCV is one of the viral proteins that interacts with viperin. NS5A interacts with the viral RNA-dependent RNA polymerase, NS5B, within the replication complex and is essential for genome replication (26,27). The reported interaction of NS5A and viperin could either be a mechanism by which viperin inhibits viral replication, or an adaptation of the virus to inhibit viperin. To investigate these possibilities we have examined the enzymatic activity of viperin in complex with NS5A and the cellular protein VAP-33, which is co-opted as part of the viral replication complex.

By reconstituting the complex between viperin, NS5A and VAP-33 in HEK293T cells, we were able to investigate how NS5A and VAP-33 alter viperin’s catalytic activity. The most significant difference in viperin activity was apparent when comparing the complex of viperin and VAP-33 and the complex of viperin, VAP-33 and NS5A, with the addition of NS5A reducing the activity of viperin by ~ 3 fold. This suggests that NS5A may have evolved, in part, to counteract the synthesis ddhCTP, thereby potentiating the infectivity HCV. The localization of these proteins to the ER membrane or lipid droplets appears to be crucial for the complex to form. This was evident from our *in vitro* studies using purified proteins, which necessitated the removal of the membrane associated domains from the proteins to facilitate their expression and purification. The truncated proteins failed to associate with each other as evident from *in vitro* co-immunoprecipitation and chemical crosslinking experiments. This observation is in agreement with previous studies on swine fever virus in which the N-terminal domain of viperin was shown to be crucial to co-localize with the replication complex at lipid droplets in HCV and interact with NS5A (28).

Although NS5A appears to inhibit viperin, our studies find support for the hypothesis that viperin exerts an antiviral effect by promoting the degradation of NS5A through the proteasomal degradation pathway. This presumably occurs by ubiquitination of NS5A. In this respect we note that viperin has been shown to activate the E3 ubiquitin ligase TRAF6 in the context of innate immune signalling. It seems plausible that viperin could act to recruit other E3 ligases that function in the endoplasmic reticulum-associated protein degradation (ERAD) pathway. Notably, this function of viperin does not appear to require the iron-sulfur cluster, as the viperinΔ3C variant was equally effective in reducing the cellular levels of NS5A. These findings are in accord with other studies showing that viperin promotes the degradation of non-structural protein NS3 in other flaviviruses through the proteasomal degradation pathway (8).

In conclusion, we find opposing effects on viral replication for the interaction between viperin and HCV NS5A *[Figure 7]*. On one hand, NS5A appears to inhibit the activity of viperin, thereby reducing the potential for ddhCTP to interfere with replication of the viral genome. On the other hand, viperin appears to decrease the expression of NS5A to limit the formation of the replication complex. These studies point to complex interplay between viral proteins and cellular proteins they co-opt for their replication.

**Figure 7:**
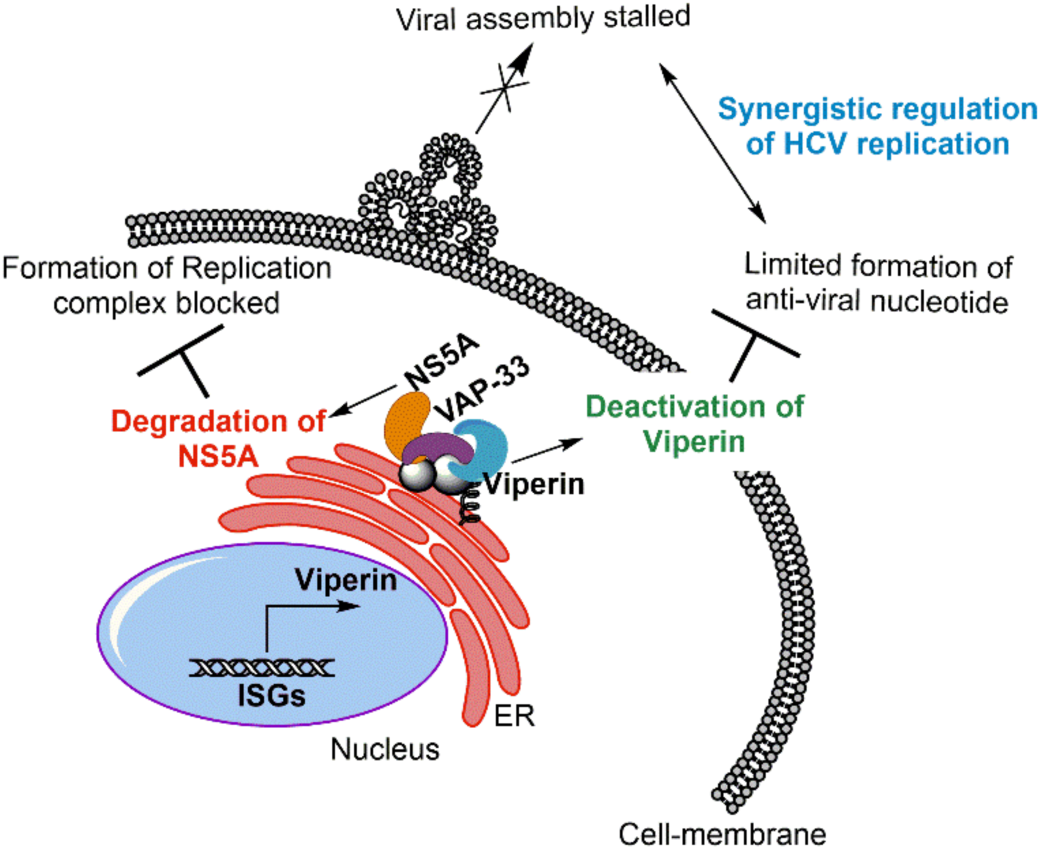
Overview of the functional interactions between NS5A and viperin. Viperin exerts anti-viral activity against Hepatitis C virus by promoting the degradation of NS5A in the replication complex. In contrast, NS5A reduces the catalytic activity of viperin, decreasing ddhCTP levels. Membrane localization of viperin, NS5A and proviral adaptor protein VAP-33 is crucial for the interactions between these proteins.

## Material and Methods

### Cell lines

The HEK293T cell line was obtained from ATCC. *E. Coli* strain BL21(DE3) was acquired from New England Biolabs.

### Plasmids

Synthetic genes encoding human viperin (GenBank accession numbers AAL50053.1) were purchased from GenScript and sub-cloned into the pcDNA3.1(+) vector (Invitrogen). The primers for the pcDNA3.1(+)-viperin construct introduced an N-terminal 3x-FLAG tag (DYKDHDGDYKDHDIDYKDDDDK) and a Kozak consensus sequence (5’-GCCAAC-3’) for downstream protein expression in eukaryotic cells. An additional construct of viperin, truncating the first 50 amino acids from the full length construct, was cloned into pcDNA3.1(+) between HindIII and EcoRI sites, using Gibson Assembly (New England Biolabs). The point mutations described herein were introduced using the QuikChange site-directed mutagenesis kit (Agilent Technologies, Mississauga, ON). The gene encoding the NS5A (genotype 1b) protein was obtained from AddGene as the pCMV-Tag1-NS5A plasmid and was sub-cloned into the pcDNA3.1(+) vector with N-terminal Myc-tag (EQKLISEEDL) using Gibson Assembly. The VAP-33 construct (hVAPc, VAP-33 residues 156-242) with N-terminal 3x-FLAG tag in pcDNA3.1(+) was a kind gift from professor Zhenghong Yuan, Fudan University, Shanghai, China(1).

For expression of vipeirn in E. coli, a codon-optimized gene encoding viperin in pET28a vector lacking the first 50 amino acids of the N-terminal amphipathic alpha helix (viperin-ΔN50, residues 50-361) was purchased from Genscript. Viperin-ΔN50 was sub-cloned into pRSFDuet-1 using the MfeI and HindIII restriction sites with a C-terminal 6x-His tag using Gibson Assembly. For expression of recombinant NS5A in E.Coli, a codon-optimized NS5A gene, lacking first 39 amino acids of N-terminal amphipathic helix (NS5A-ΔN39, residues 39-447), housed in pET52b(+) was purchased from Genscript. NS5A-ΔN39 was sub-cloned downstream of maltose binding protein (MBP) in pMALc5x between the NdeI and HindIII restriction sites, using Gibson Assembly. This added a tobacco etch virus (TEV)-cleavable MBP tag to the N-terminus with a non-cleavable 6x-His tag (encoded in the downstream primer) on the C-terminus of NS5A. The pMALc5x and pRSFDuet-1 vectors were obtained from GenScript. All sequences were confirmed at the University of Michigan Biomedical Research Core Facilities Advanced Genomics Core. Recombinant human VAP-33 protein (VAP-33-ΔC20, residues 1-227) with an N-terminal 6x-His tag was purchased from IBL America (catalog number IBATGP0471).

### Cell Culture and Transfection

The pcDNA3.1(+) encoded constructs were overexpressed in HEK293T cells (cultivated in DMEM supplemented with 10% FBS and 1% antibiotics) through transient transfection using polyethyleneimine transfecting agent. Co-transfection of two or more DNA constructs, allowing co-expression of viperin and its target proteins within the HEK293T cells, was performed using the same transfection protocol. Briefly, 20 µg of the respective plasmid DNA construct was mixed with polyethyleneimine at a 1:2 ratio, incubated at room temperature for 10 minutes, and then added to HEK293T cells at 50-60% confluence on a 100-mm dish. The transfected cells were then grown for up to 36 hours, gently pelleted, and stored at −80 °C until use.

### Antibodies

Rabbit polyclonal RSAD2/ viperin antibody (11833-1-AP) and mouse monoclonal viperin (MABF106) were obtained from Protein Tech and EMD Millipore respectively. Mouse monoclonal anti-NS5A antibody was purchased from Virogen (256-A). Goat anti-rabbit (170–6515) and anti-mouse (626520) Ig secondary antibodies were purchased from BioRad and Life Technologies respectively. Rabbit polyclonal GAPDH (10494-1-AP) was purchased from Proteintech and mouse monoclonal GAPDH antibody (6C5) was obtained from EMD Millipore. Mouse monoclonal anti-FLAG® M2 antibody (F1804) and 6x-His tag monoclonal antibody (MA1-135) were purchased from Sigma-Aldrich and ThermoFisher Scientific. Goat anti-mouse IgG (H+L)-HRP conjugate secondary antibody (catalog number 626520) and goat anti-rabbit IgG (H+L)-HRP conjugate secondary antibody (catalog number 1706515) were purchased from ThermoFisher Technologies and Bio-Rad, respectively.

### Reagents

S-(5ʹ-Adenosyl)-L-methionine *p*-toluenesulfonate salt (≥80% (HPLC), 25 MG-A2408), 5’-deoxyadenosine (D1771-25MG) was purchased from Sigma-Aldrich. Sodium Hydrosulfite (7775-14-6, 100g) and DTT (DSD11000-25 MG) were purchased from Fisher Scientific and DOT Scientific Inc. respectively. MG-132 10 mM solution in DMSO, 1 mL (A11043) was purchased from AdooQ Bioscience. Nucleotide substrates CTP (Cytidine 5’-triphosphate, disodium salt hydrate, 95%) were purchased from Acros Organics-226225000. Pierce™ protein A/G plus agarose resin and control agarose resin (Pierce classic IP kit 26146) were purchased from ThermoFisher Scientific. Transfection Grade Linear Polyethylenimine Hydrochloride (MW 40,000) (Catalog No. 24765-1) was purchased from Polysciences, Inc. for DNA transfection in HEK293T cells. For *in vitro* chemical crosslinking experiment, bis(sulfosuccinimidyl)suberate or (BS^3^) (catalog number 21580) was purchased from Thermo Fisher Scientific.

### Immunoblotting

Cells were lysed in lysis buffer (20 mM Tris, pH 7.5, 1% NP-40, 150 mM NaCl, 1 mM EDTA, 1 mM Na_3_VO_4_, and 10 mM β-glycerophosphate with SIGMAFAST™ Protease Inhibitor Tablets, S8830; Sigma). Supernatants of lysates were collected and mixed with 4x Laemmli sample buffer and 2-mercaptoethanol. The total amount of protein in lysates was determined by DC™ Protein Assay (5000116, Biorad). The supernatants were separated on 10% SDS-PAGE gels and transferred to PVDF membrane. Membranes were blocked for 1 h at room temperature in TBST buffer (20 mM Tris, pH 7.5, 137 mM NaCl, and 0.1% Tween 20) containing 5% nonfat dry milk, followed by overnight incubation at 4°C in TBST buffer containing 5% nonfat dry milk and the appropriate primary antibody. Membranes were washed three times in TBST and then incubated for 1 h at room temperature with the secondary IgG-coupled horseradish peroxidase antibody. Primary antibodies were used at the following dilutions: viperin constructs - rabbit polyclonal diluted 1:2500, GAPDH - rabbit polyclonal diluted 1:5000, NS5A - mouse monoclonal diluted 1:5000, anti-FLAG - mouse monoclonal diluted 1:3000, and anti-His - mouse monoclonal diluted 1:3000. Secondary antibodies were used at the following dilutions: goat anti-mouse diluted 1:5000 and goat anti-rabbit diluted 1:5000. The membranes were washed three times with TBST, and the signals were visualized with enhanced chemilluminescence reagent. Band intensities quantified using a Bio-Rad ChemiDoc Touch imaging system. Integrated density measurements were done using ImageJ software. Quantitative measurements of protein expression levels reported here represent the average of at least three independent biological replicates.

### Co-immunoprecipitation

Cells expressing viperin/ viperinΔ3C/viperin-ΔN50, NS5A and hVAPc from 100 mm tissue culture plate were rinsed twice with ice-cold PBS, harvested in lysis buffer (20 mM Tris, pH 7.5, 1% NP-40, 150 mM NaCl, 1 mM EDTA, 1 mM Na_3_VO_4_, and 10 mM β-glycerophosphate with EDTA-free protease inhibitor cocktail from Sigma), incubated on ice for 20 minutes and briefly sonicated. Lysates were collected by centrifugation at 13,000 × g for 15 minutes at 4°C and pre-cleared with 20 µl Pierce Control Agarose Resin. Lysates containing bait protein viperin/ viperinΔ3C/TN50-vip were incubated with rabbit polyclonal anti-viperin antibody for 1 h at 4°C with end-to-end rotation. The protein-antibody-complexes were incubated with pre-equilibrated Pierce protein A/G plus agarose resin at the ratio of suspension to packed gel volume 4:1 for 1 h at 4°C by end-to-end rotation. Lysates containing prey protein NS5A and hVAPc were incubated with bait proteins at 1:1:1 total protein ratio for 30 minutes at 4°C. Flow-through was collected by gravity flow using Pierce gravity – flow columns and washed three times with wash buffer (20 mM Tris, pH 7.5, 1% NP-40, 150 mM NaCl, 1 mM EDTA, 1 mM Na_3_VO_4_, and 10 mM β-glycerophosphate). Immuno-complexes were eluted by boiling in 1x SDS-PAGE sample buffer and immunoblotted with the appropriate antibodies.

### Immunofluorescence

HEK293T cells were grown to 30% confluence on poly-L-lysine-coated coverslips and then transiently transfected with wild-type viperin, viperinΔ3C, NS5A and hVAPc plasmid DNA. After 36 hours of transfection, the cells were fixed with 1% paraformaldehyde, permeabilized with 0.05% Triton X-100 dissolved in PBS, and washed three times with PBS containing 0.1% Tween20. The fixed cells were stained with the appropriate antibodies after blocking with 1% FBS in PBS. Primary antibodies were diluted in PBS containing, 1% FBS and stained with rabbit polyclonal anti-viperin (Proteintech) and mouse monoclonal anti-NS5A (Virogen) antibodies at 1:500 dilution. After incubation at 4°C overnight, the coverslips were washed with PBS containing 0.1% Tween20 and treated with Alexa Fluor 647-conjugated goat anti-mouse (Life Technologies) and Alexa Fluor 488-conjugated goat anti-rabbit (Abcam) secondary antibodies at a dilution of 1:500 at room temperature for 2 hours. The coverslips were washed three times with PBS containing 0.1% Tween20 and mounted in ProLong™ Gold Antifade Mountant (Molecular Probes). Images were acquired with an Olympus IX81 microscope with 60x objective. The images were processed with ImageJ software.

### Reductive SAM assay of viperin

HEK293T cells transfected with viperin, and/or NS5A and hVAPc were harvested from one 100-mm diameter tissue culture plate each, re-suspended in 500 µl of anoxic Tris-buffered saline (50 mM Tris-Cl, pH 7.6, 150 mM NaCl) containing 1% Triton X-100, sonicated within an anaerobic glovebox (Coy Chamber), and centrifuged at 14,000 g for 10 min. Dithiothreitol (DTT; 5 mM) and dithionite (5 mM) were added to the cell lysate together with CTP (300 μM). The assay mixture was incubated at room temperature for 30 min prior to starting the reaction by the addition of SAM (200 µM). The assay was incubated for 60 min at room temperature, after which the reaction stopped by heating at 95 °C for 10 min. The solution was chilled to 4 °C, and the precipitated proteins were removed by centrifugation at 14,000 rpm for 25 min. The supernatant was then extracted with acetonitrile. Samples were analysed in triplicate by UPLC-tandem mass spectrometry(29).

### NS5A-ΔN39 in vitro expression and purification

MBP_NS5A-ΔN39 was transformed into BL21(DE3) chemically competent cells and plated on ampicillin (AMP) supplemented LB-agar plates. The following day, a single colony was picked and added to 2-25mL fractions of AMP-supplemented LB media and shaken overnight at 37°C. The seed cultures were added to 2L of 2xYT media supplemented with AMP and 1% glycerol. The culture was shaken for approximately 3 hours, until mid-log was achieved (O.D._600_ ≈ 0.8), then cold shocked by incubation at 4°C for one hour. 0.5 mM IPTG was added and the culture was grown at 16°C overnight. The cells were centrifuged and the pellet was harvested and stored at −80°C.

The cell pellet was re-suspended at a ratio of 30 mL/ 2L culture in NS5A lysis buffer (30)(50 mM Tris pH 8.0, 200 mM NaCl, 10 mM imidazole, 5% glycerol, 2 mM L-Cysteine – HCl, 2 mM 2-mercaptoethanol) supplemented with 1 cOmplete^TM^ EDTA-free protease inhibitor cocktail tablet. The cell suspension was lysed on ice via sonication at amplitude 8 in 20 second intervals for a total process time of 4 minutes. The lysate was centrifuged for 45 min at 12,000 RPM. 1 mL of 100% Ni-NTA resin (pre-equilibrated with lysis buffer) was added to the supernatant and incubated at 4°C for 2 hr. The mixture was added to a 20 mL fritted column and the flow-through was collected and set aside. The column was washed with 30 column volumes of lysis buffer supplemented with 25 mM imidazole. The protein was eluted from the Ni-NTA using three - 10 column volume washes of lysis buffer supplemented with 100, 200, or 300 mM imidazole. The lysate, pellet, supernatant, flow through, wash, and all elutions were analysed via reducing sodium dodecyl sulfate – polyacrylamide gel electrophoresis (SDS-PAGE).

### NS5A-ΔN39 TEV cleavage and size exclusion chromatography

Once presence and purity of the protein was confirmed, the appropriate elution fraction was subjected to TEV cleavage. The reaction contained approximately 10-15 mg of pure MBP_NS5A-ΔN39, 50 μg TEV, and 3 mM DTT, was carried out overnight (18-20 hours) at 4°C, and the cleaved protein was purified over a pre-packed 5 mL MBP-trap column (from GE). The reaction was circulated over the column via peristaltic pump at 0.5 mL/min three times and the flow through, which contained cleaved NS5A, was collected after a fourth pass. The column was washed with 10 column volumes of NS5A storage buffer (50 mM Tris pH 8.0, 150 mM NaCl, 10% glycerol). Any un-cleaved protein was eluted with 5 column volumes of NS5A storage buffer supplemented with 10 mM maltose.

Next, a size exclusion step was added in order to remove any soluble aggregate and additional protein contaminants not removed by the post-TEV cleavage purification. Cleaved NS5A-ΔN39 was concentrated down to approximately 3 mL and injected onto a pre-equilibrated prep-grade Superdex-200 size exclusion column. The column was washed with NS5A storage buffer without glycerol at 1 mL/min. Elution fractions were compared against a 15-600 kDa protein standard (Millipore Sigma) and analysed via SDS-PAGE for purity and presence of soluble, cleaved NS5A-ΔN39. These fractions were concentrated and buffer exchanged using a PD-10 desalting column into NS5A storage buffer. This was concentrated further to 500 μL (final concentration ~ 38 μM), aliquoted, and flash frozen for future use. Protein concentration was estimated via Abs_280_ and confirmed with gel quantification.

### Viperin-ΔN50 in vitro expression and purification

Viperin-ΔN50 was expressed in the same manner as described above with small differences. pRSFDuet-1 is kanamycin (KAN) resistant, so all antibiotic selection steps were done with KAN. Once the culture entered mid-log phase, it was equilibrated to 18°C after which 0.2 mM Na_2_S · 9H_2_0 was added. After 20 additional minutes of equilibration, 0.2 mM FeCl_3_ and 0.1 mM IPTG were added and the culture was shaken overnight at 16°C. Cells were harvested by centrifugation the following day and stored at −80°C.

The purification was carried out in the same manner as above, with small differences. All purification steps were done in an anaerobic environment (Coy chamber) using anoxic (nitrogen flushed) buffers (viperin-ΔN50 lysis buffer contained 50 mM HEPES pH 7.5, 300mM NaCl, and 5% glycerol). Once the lysate had been cleared via centrifugation it was added to a pre-packed 5 mL His-Trap column (from GE) via peristaltic pump. The solution was flowed over the column 3x at 0.5 mL/min and the flow through was collected on the fourth pass. The column was washed with 10 column volumes of viperin-ΔN50 lysis buffer supplemented with 50mM imidazole. viperin-ΔN50 was eluted with three 15 mL washes of lysis buffer supplemented with 200 mM imidazole. Protein presence and purity was analysed in the same manner as above. Once confirmed, the appropriate elution fraction was concentrated, buffer exchanged into viperin storage buffer (50mM Tris pH 8.0, 150 mM NaCl, 10% glycerol) with a PD-10 desalting column, and stored on ice before reconstitution.

### Reconstitution of viperin-ΔN50 Fe_4_S_4_ cluster

The reconstitution of the Fe_4_S_4_ cluster in the purified viperin-ΔN50 was performed in an anaerobic environment (Coy chamber, O_2_ levels kept below 20 ppm)(15). Purified protein was incubated with 5 mM DTT for 20 min on cold beads. The stock solution of 0.1 M FeCl_3_ and 0.1 M Na_2_S was added drop-wise, incubating for 10 min after each addition until the protein turned dark brown in color. A 10 mL PD-10 desalting column was equilibrated with 2 column volumes of viperin storage buffer. The reconstituted protein (2.5 mL) was then added to the equilibrated PD-10 column and eluted with 3.5 mL of viperin storage buffer. The elution was then concentrated again using vivapore concentrators to the final volume of 1 mL. The concentrated protein (final concentration~ 68 μM) was aliquoted, flash frozen in liquid nitrogen, and stored at - 80°C.

### Co-immunoprecipitation of recombinant NS5A and VAP-33 using recombinant viperin

All steps were performed inside anaerobic Coy chamber. Recombinant human viperin-ΔN50, purified from BL21 (DE3) *E. Coli*, was incubated with rabbit polyclonal anti-viperin antibody for 1h at 4°C, at final concentration 0.5 μM, in the presence of dithiothritol (5 mM) and CTP (300 μM). The protein-antibody complex was incubated with pre-equilibrated protein A/G beads (pre-blocked with 2mg/ml Bovine Serum albumin protein overnight at 4°C) for 1h at 4°C. Recombinant prey proteins NS5A-ΔN39 and VAP-33-ΔC20 were incubated with viperin-antibody-bead-complex at a final concentration of 1µM for 30 minutes at 4°C. Flow-through was collected using Pierce gravity columns and washed three times with 50 mM Tris, pH 7.5, 150 mM NaCl, 10% glycerol, 1.0% Tween-20. Immuno-complexes were eluted by boiling in 1x SDS-PAGE sample buffer outside Coy chamber and immunoblotted with appropriate antibodies.

### In vitro SAM cleavage assay

The reaction containing purified 5 uM viperin-ΔN50, 7.5 uM NS5A-ΔN39 or VAP-33-ΔC20 or both in 50 mM tris-HCl pH 8.0, 150 mM NaCl, 10% glycerol, 5 mM DTT, 100 uM L-tryptophan (internal standard), and 300 uM CTP were incubated on cold beads for 2 hours under anaerobic conditions to allow for complex formation. The reactions were then incubated at 37°C for 5 minutes after the addition of 5 mM dithionite. Finally, 200 uM SAM was added to each reaction and 20 uL aliquots were taken at various time intervals (0, 2, 4, 6, 8, and 10 minutes). Each sample was quenched with 20 uL of 50 mM H_2_SO_4_ solution. The samples were centrifuged at 14,000 rpm for 10 minutes to precipitate protein before loading it on to a Vydac 201TP 10 µm C18 column (250 × 4.6 mm, 10 µm particle size). Buffer A was 0.01% TFA in DI water and buffer B was 0.01% TFA in acetonitrile. The flow rate was 1.0 ml/min and the following gradient was applied: 0% B for 0.01 minutes, 0-5% B from 0.01-5.01 minutes, 5% B from 5.01-5.31 minutes, 5-75% B from 5.31-25.31 minutes, 75% B from 25.31-26.31 minutes, 75-100% B from 26.31-27.01 minutes, 100% B from 27.01-32.01 minutes, 100-0% B from 32.01-32.31 minutes, 0% B from 32.31-36.31 minutes. The internal standard peak (L-tryptophan) was observed at 13.25 minutes and the 5’deoxyadenosine (5’dA) peak was observed at 10.10 minutes. The peaks were integrated using LCSolution software.

## Acknowledgements

This research was supported by the National Institutes of Health, grant GM 093088 to E.N.G.M.

---

**Figure SI.1:**
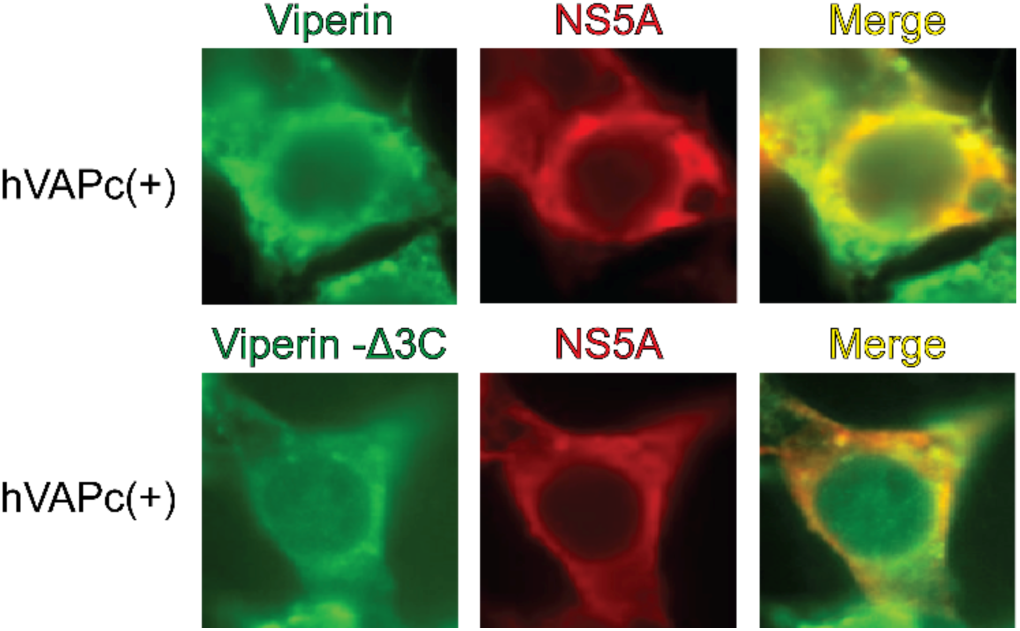
NS5A co-localizes with viperin or viperin-Δ3C at endoplasmic-reticulum. Immunofluorescence microscopy of HEK293T cells co-transfected with viperin (*upper panel*) or viperin-Δ3C (*lower panel*) and NS5A, in presence of hVAPc. The cells were immobilized 30 hours post-transfection and stained for viperin (green) and NS5A (red). Both viperin and viperin-Δ3C co-localize (yellow in merged images) with NS5A.

**Figure SI.2:**
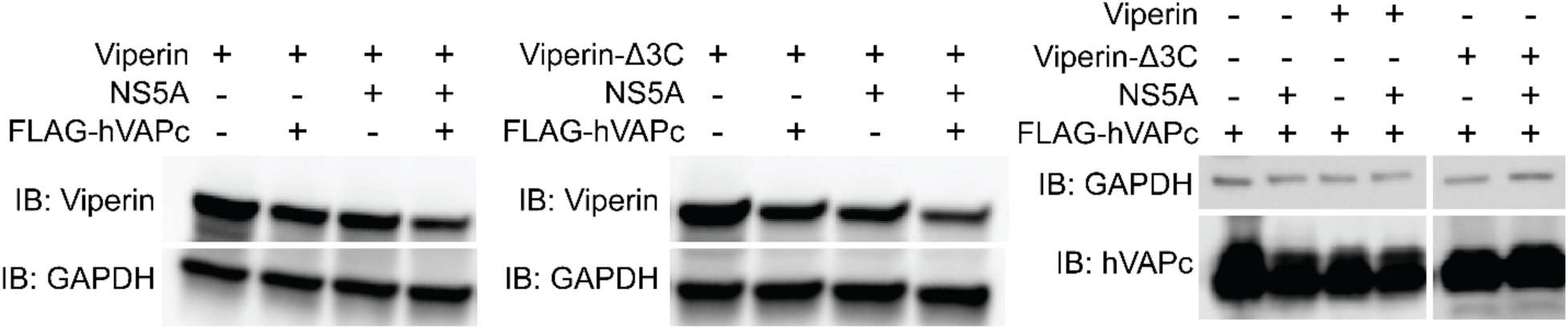
Expression level of viperin, viperin-Δ3C and hVAPc. HEK293T cells were transfected with genes of NS5A, FLAG-hVAPc and viperin or viperin-Δ3C and immunoblotted for viperin, viperin-Δ3C, hVAPc and GAPDH (loading control) 30 hours post-transfection. (a) Viperin and viperin-Δ3C, showed reduced expression when co-transfected with NS5A and hVAPc. (b) The expression level of hVAPc remained the same, regardless the presence of viperin and NS5A.

**Figure SI.3:**
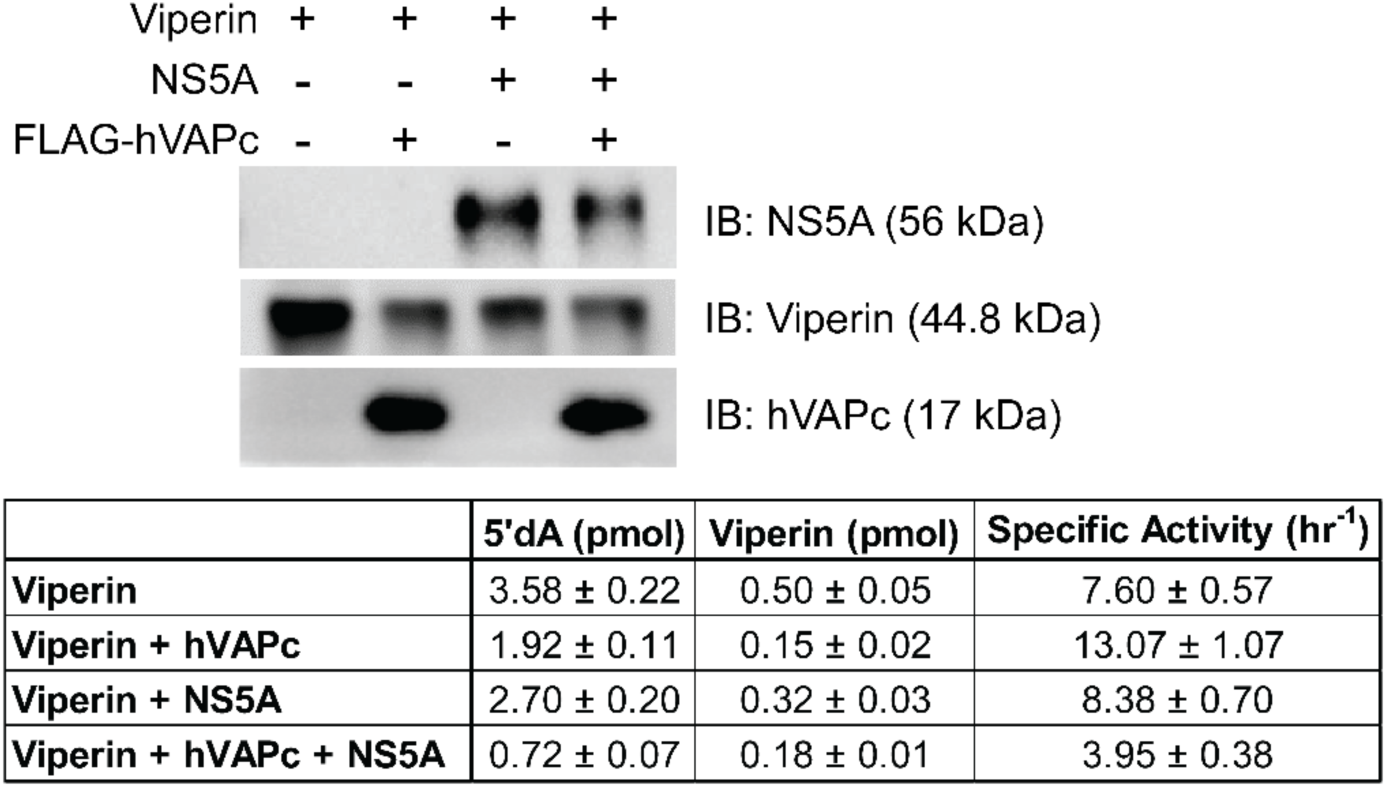
Co-expression of NS5A and hVAPc inhibits reductive SAM cleavage activity of Viperin in HEK293T cell-lysates (contd.). (a) Immunoblotting of vipeirn, NS5A and hVAPc present in the samples. (b) The relative activity was of viperin was determined by the ratio of 5’-deoxyadenosine produced to amount of viperin present in the sample per hour.

**Figure SI.4:**
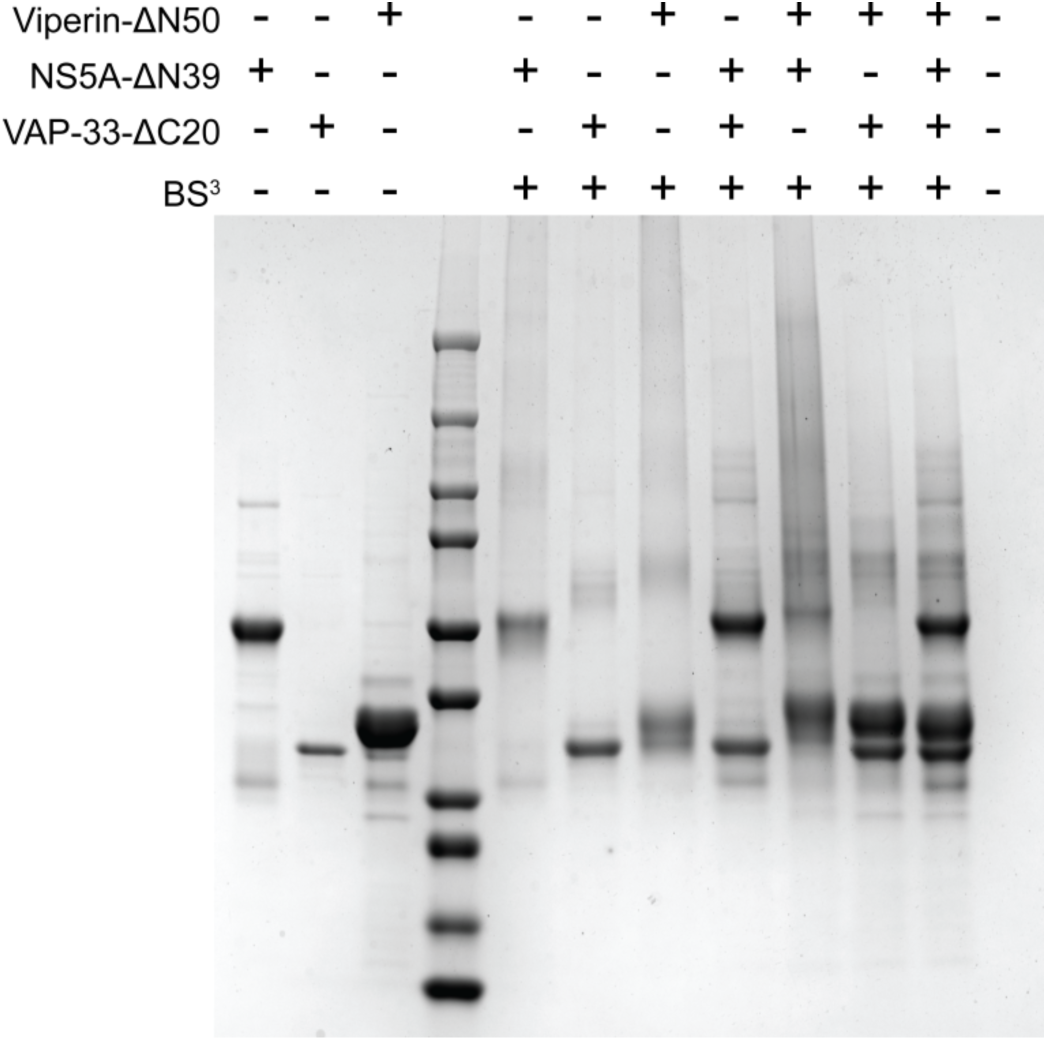
Non-specific oligomerization of recombinant viperin, NS5A and VAP33 in vitro after chemical crosslinking assay. Purified viperin (viperin-ΔN50), NS5A (NS5A-Δ39) and VAP-33 (VAP-33-ΔC20) were incubated with each other or by itself prior to the addition of chemical crosslinker BS^3^. The three proteins individually formed dimers and showed non-specific oligomerization in the presence other partner protein. The protein complex among these three proteins was not observed.

